# Elevated peripheral and nervous system inflammation is associated with decreased short-chain fatty acid levels in Zika-virus infected macaques

**DOI:** 10.1101/2023.07.25.550459

**Authors:** Charlene J. Miller, Jennifer A. Manuzak, Andrew T. Gustin, Christopher M. Basting, Ryan K. Cheu, Ty A. Schroeder, Adrian Velez, Connor B. Driscoll, Jennifer Tisoncik-Go, Luca Schifanella, Tiffany Hensley-McBain, Claudya A. Evandy, Elise A. Smith, Debbie Bratt, Jeremy Smedley, Megan A. O’Connor, Deborah H. Fuller, Dan H. Barouch, Michael Gale, Nichole R. Klatt

## Abstract

Zika virus (ZIKV) infection of central nervous system (CNS) tissue is associated with CNS inflammation, which contributes to ZIKV pathology. Similarly, ZIKV infection has been associated with increased vaginal and rectal mucosal inflammation. As mucosal dysfunction may contribute to elevated systemic inflammation, ZIKV-induced mucosal alterations could potentiate CNS disruptions, leading to ZIKV pathogenesis. However, the potential link between mucosal dysfunction, CNS inflammation and the underlying mechanisms causing these disruptions in ZIKV infection has not been well described. Here, we assessed plasma and CSF indicators of inflammation, including neopterin, tryptophan, kynurenine and serotonin by liquid chromatography tandem mass spectrometry. We observed significant increases in neopterin formation, tryptophan catabolism and serotonin levels in the plasma and CSF of ZIKV-infected pigtail macaques (PTM), rhesus macaques (RM) and in the plasma of ZIKV-infected humans. We next examined whether ZIKV infection resulted in microbial translocation across mucosal surfaces by evaluating plasma and cerebrospinal fluid (CSF) levels of soluble CD14 (sCD14) and lipopolysaccharide-binding protein (LBP) by enzyme-linked immunosorbent assay (ELISA). Increased sCD14 was observed in the CSF of PTM and rhesus macaque (RM), while increased LBP was observed in pigtail macaque (PTM) plasma. Finally, to examine whether ZIKV-induced microbial dysbiosis could underlie increased microbial translocation and inflammation, we characterized intestinal microbial communities by 16s rRNA gene sequencing and microbial functional changes by quantifying short-chain fatty acid (SCFA) concentrations by gas chromatography mass spectrometry. We observed that although ZIKV infection of PTM did not result in significant taxonomic shifts in microbial communities, there were significant reductions in SCFA levels. Loss of microbial function in ZIKV infection could cause decreased intestinal integrity, thereby contributing to elevated microbial translocation and systemic and CNS inflammation, providing a possible mechanism underlying ZIKV pathogenesis. Further, this may represent a mechanism underlying inflammation and pathogenesis in other diseases.

**Author Summary:** Zika virus (ZIKV) can be transmitted to humans via the bite of an infected mosquito or between humans during sexual intercourse, typically resulting in mild symptoms, which has been linked to elevated inflammation in the CNS and the development of more serious conditions, including severe neurological syndromes. Previous studies have observed that ZIKV infection is associated with increased mucosal dysfunction, including elevated inflammation in rectal and vaginal mucosal tissue. However, the mechanism of ZIKV-induced mucosal dysfunction may contribute to systemic and CNS inflammation has not been previously investigated. Here, we used the non-human primate (NHP) model and clinical specimens from ZIKV-infected humans to examine markers of systemic and CNS inflammation and microbial translocation. We observed elevated markers indicative of microbial translocation and inflammation in the CNS of ZIKV-infected macaques and humans. A potential association with mucosal dysfunction in ZIKV infection is shifts in microbial dysbiosis. We also observed that there were no significant overall taxonomic shifts in microbial communities, but a reduction of bacterial-derived short-chain fatty acid (SCFA) levels. Finally, we observed that the decrease in SCFA levels significantly negatively correlated with the elevated peripheral and CNS inflammatory markers, suggesting a link between ZIKV-driven disease pathology and microbial function. Taken together, our study provides new insight into a previously unconsidered mechanism underlying ZIKV pathogenesis.

## Introduction

As has been acutely evident by the Covid-19 pandemic, viral infections are a major health problem, and understanding viral-induced effects on human health is critical. Zika virus (ZIKV) is an arbovirus of the Flaviviridae family that threatens the health of millions globally and was the cause of a major pandemic in 2015-2016. ZIKV infection typically results in mild symptoms including rash, fever and conjunctivitis in adults, but it has occasionally been linked to more severe conditions, including Guillain-Barré syndrome (GBS) and Zika congenital syndrome (1). Previous work assessing a Brazilian cohort of ZIKV-infected human adults demonstrated an increased incidence of serious neurological syndromes in infected individuals, such as demyelinating GBS, axonal GBS, encephalitis and transverse myelitis (2). Serologic studies have detected Zika viral infections among humans globally, most recently in the Americas and Caribbean (3).

ZIKV has been detected in central nervous system (CNS) tissue (2,4–8), which likely causes ZIKV-associated neuropathy and neuroinflammation (6,9–11). However, additional mechanisms may drive CNS inflammation in the context of ZIKV infection. For example, inflammatory monocyte and macrophage accumulation in the brain has been observed during viral infections, including other Flavivirus infections like West Nile Virus (12) and Japanese encephalitis virus (13). Accumulation of activated monocytes/macrophages in the CNS during ZIKV infection could result in elevated production of neopterin, a biomarker of immune activation produced by monocytes and macrophages in response to interferon stimulation (14,15). Indeed, elevated neopterin levels have been observed in the cerebrospinal fluid (CSF) of individuals with viral or bacterial CNS infections (16). Additionally, monocyte activation via IFN-γ stimulation or recognition of pathogen associated molecular patterns (PAMPs) via pattern recognition receptors (PRRs) could induce the gene expression and enzymatic activity of indoleamine 2,3-dioxygenase 1 (IDO-1), the first enzyme involved in the kynurenine pathway of tryptophan catabolism (17–20). Altered tryptophan catabolism and kynurenine levels have been observed in many different viral infections (18), as well as in CNS disorders like AIDS-dementia complex and Huntington’s disease (21–23). Thus, innate immune cell inflammatory and metabolic responses could contribute to CNS inflammation in the context of ZIKV infection. However, the impact of ZIKV infection on these processes and their role in ZIKV pathogenesis has not yet been investigated.

In addition to the detrimental effects of ZIKV infection in the CNS, previous work has shown that ZIKV can replicate in mucosal compartments, including the intestinal and vaginal mucosa (24,25). Elevated intestinal mucosal dysfunction due to ZIKV infection (24) could potentiate systemic and eventually CNS inflammation. For example, disruptions in mucosal barrier integrity during ZIKV infection could allow for microbial translocation from the lumen of the gastrointestinal (GI) tract into the periphery and CNS. Increased exposure to these translocated microbial products could drive CNS inflammation, potentially by inducing neopterin production or tryptophan catabolism in monocytes. Additionally, GI microbial dysbiosis, defined as imbalances in microbial communities, has been observed in chronic viral infections, including HIV (26–28) and viral hepatitis (29). As the intestinal microbiome plays an important role in the maintenance of intestinal homeostasis (30), shifts in microbial taxonomy or functionality subsequent to ZIKV infection may provide a potential mechanism for elevated microbial translocation, increased systemic inflammation and risk for negative neurological outcomes. However, no studies as yet have investigated the occurrence and potential links between ZIKV-induced mucosal disruptions, microbial translocation and dysbiosis, and peripheral and CNS inflammation.

Macaque models of HIV, influenza, Ebola virus and others, display infection kinetics and pathology like human infections, enabling the detailed study of these diseases (31). Given the difficulty in identifying and sampling acute ZIKV infection in humans, the macaque model is similarly invaluable for studying the earliest stages of ZIKV pathogenesis. Recent studies in rhesus macaques have detected persistence of ZIKV in multiple tissues as late as 35 days post-infection (dpi) (32). Additionally, ZIKV infection elicits a robust immune response in rhesus macaques that includes ZIKV-specific T cell and neutralizing antibody (Nab) responses that confer protection against reinfection (33). Finally, our group has previously shown that ZIKV infection in pigtail macaques results in a rapid innate immune response in the periphery and recruitment of innate immune cells to the intestinal mucosa (24).

Given the limited knowledge regarding whether ZIKV-associated mucosal dysfunction could drive CNS inflammation and overall ZIKV pathogenesis, here we sought to evaluate the link between intestinal mucosal alterations and peripheral and CNS immune disruption in the context of ZIKV infection. To do this, we used the pigtail macaque (PTM) model to evaluate associations between the level of microbial translocation and systemic and CNS innate immune activation and metabolic activity during ZIKV infection. Additionally, we explored ZIKV-induced alterations in intestinal microbial community structure and function as a potential mechanism underlying systemic and CNS inflammation. Finally, to support our PTM findings, we performed similar assessments using samples from ZIKV-infected rhesus macaques (RM) and humans.

## Results

### Infection and sampling of macaques with ZIKV

Pigtailed macaques (PTM; n=8) were infected with the Brazilian Zika isolate (Brazil_2015_MG, GenBank: KX811222.1). ZIKV inoculation and sample collection was completed in two separate, non-overlapping cohorts of 4 animals each (Cohort 1: n=4 female PTM; Cohort 2: n=4 male PTM). ZIKV inoculum dose was based on the feeding behavior of the ZIKV mosquito vector, *Aedes aegypti*, which deposits virus multiple times during skin probing prior to accessing a capillary for acquisition of the bloodmeal (34). Animals did not exhibit overt clinical symptoms subsequent to ZIKV inoculation. However, we observed that anemia occurred in the females, which could also be associated with menses. Additionally, macroscopic mucosal shedding was observed in the gastrointestinal tract during endoscopic procedures post-infection. As previously shown, PTMs exhibited plasma viremia 3 dpi, but viral loads dropped to undetectable levels by 7 dpi (24). Blood, stool, rectal swabs and cerebral spinal fluid (CSF) were collected pre- and post-infection with ZIKV. The data presented in the current study were generated from cryopreserved samples after all sample collection for both cohorts had been completed. Of note, gastrointestinal (GI) sampling (rectum and colon biopsies collected via endoscopy) was also performed in this PTM study, and data generated using these tissues have been previously published (24).

In addition, plasma and CSF samples were collected from ZIKV-infected rhesus macaques (RM) enrolled in a previously completed study (4) and were used in a collaborative effort to supplement findings in the PTM model. The RMs were infected with a lower viral dose of the Brazil/AKV2015 isolate and as previously shown, exhibited lower initial viral loads that were measurable until 14 dpi (4).

### Increased markers of inflammation in CSF and plasma during ZIKV infection

Neopterin is produced by monocytes and macrophages in response to IFN-y stimulation (14,15) and is commonly used as a diagnostic marker for generalized immune activation in the context of a wide variety of infections, including bacterial, viral, and parasitic diseases (14). Elevated levels of neopterin have been observed in the CNS of individuals in many conditions, among them HIV infection (35) and CNS lymphoma (36). To determine whether ZIKV infection resulted in elevated neopterin levels in the CNS, we quantified CSF levels of neopterin in ZIKV-infected PTMs and RMs. In PTMs, a significant increase of neopterin was detected in CSF at 7 dpi, which was sustained until 14 dpi (*p*=0.0097 and 0.0075, respectively), returning to baseline by 21 dpi (Fig 1A). In the CSF of RMs, a significant increase in neopterin was detected as early as 2 dpi (*p*=0.0002), which was maintained at 7 dpi and 14 dpi (*p*=0.0009 and 0.0003, respectively), before returning to baseline at 35 dpi (Fig 1B).

**Figure 1.**
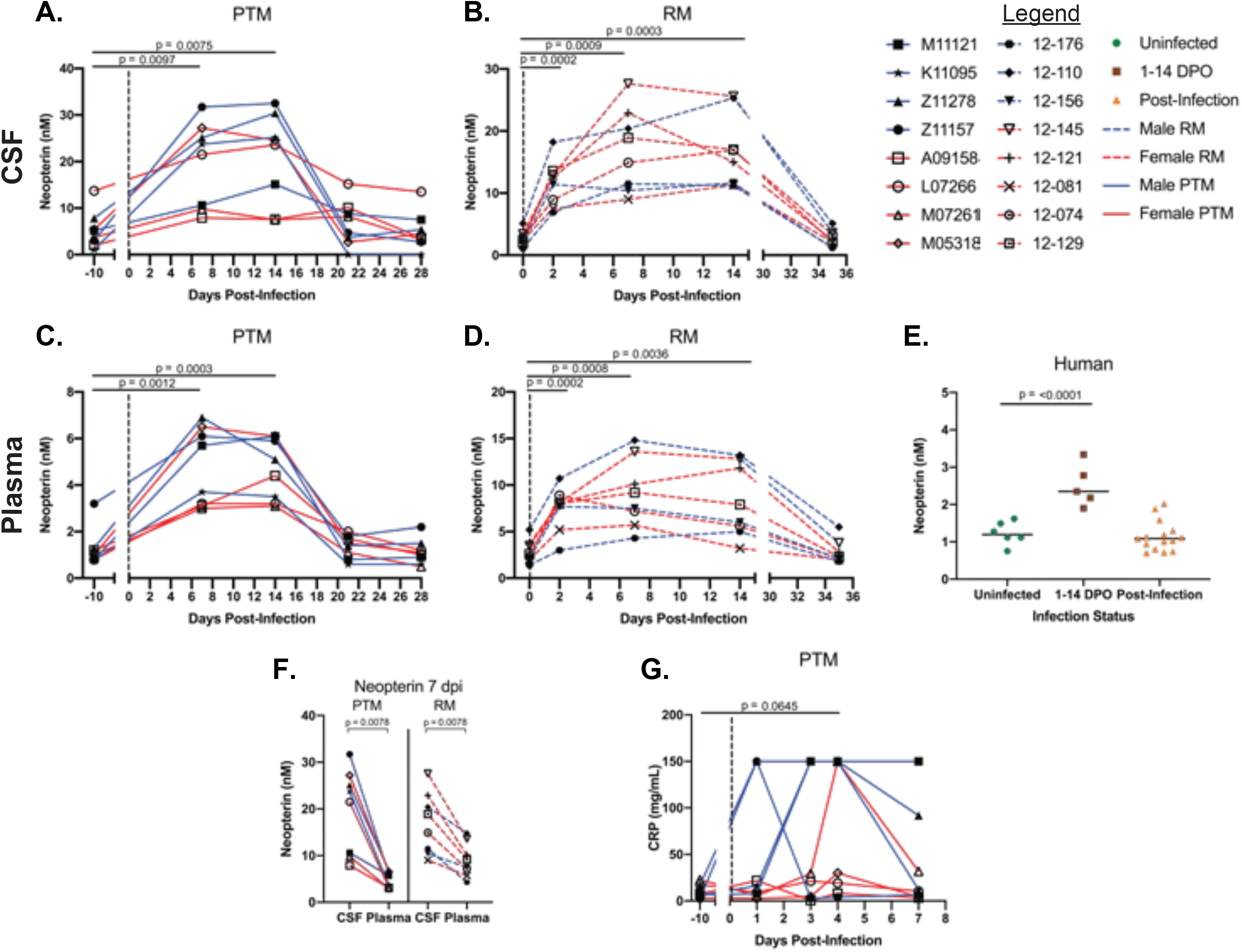
Increased levels of peripheral and CNS inflammatory markers during ZIKV infection. Liquid chromatography tandem mass spectrometry (LC-MS/MS) was used to detect neopterin concentrations in the CSF of PTM and RM and in the plasma of PTM, RM and humans throughout ZIKV infection. Additionally, CRP was measured in the plasma of PTM during acute ZIKV infection by ELISA. (A-E) Neopterin concentrations in the CSF of PTM (A), CSF of RM (B), plasma of PTM (C), plasma of RM (D) and serum of humans (E) infected with ZIKV. (F) Neopterin concentrations in CSF was compared to plasma neopterin concentrations at 7 dpi in PTM and RM. (G) Levels of CRP in plasma of ZIKV-infected PTM. In panels A-D and F-G, each animal is represented by a different symbol. Solid or dashed lines connect data from the same animal. Solid blue lines indicate male PTMs; solid red lines indicate female PTMs; dashed blue lines indicate male RMs, dashed red lines indicate female RMs. Vertical dotted lines indicate the time point at which animals were inoculated with ZIKV. Statistical significance between the pre-infection baseline and post-ZIKV infection time points was calculated using a repeated measures ANOVA, with the Geisser-Greenhouse correction and a Dunnett’s multiple comparisons post-test. In panel E, uninfected humans are indicated by green circles, humans experiencing ZIKV symptoms from 1-14 days post-onset (DPO) of ZIKV symptoms are indicated by brown squares and humans post-ZIKV infection (15-156 DPO) are indicated by orange triangles. Black bars indicate median values. Statistical significance between human sample timepoints was evaluated using a one-way ANOVA with a Dunnett’s multiple comparisons post-test. In all plots, horizontal bars with p-values indicate time points that were statistically significant from each other.

We next evaluated whether ZIKV infection resulted in elevated neopterin production in the periphery of PTMs and RMs, and we supplemented these observations with assessments of peripheral neopterin levels in ZIKV-infected humans, using serum obtained from the Global Virus Network (GVN) Zika serum bank. In PTMs, a significant increase in neopterin was detected 7 dpi and remained elevated at 14 dpi (*p*=0.0012 and 0.0003, respectively), returning to baseline by 21 dpi (Fig 1C). In the plasma of RMs, a significant increase of neopterin was detected as early as 2 dpi (*p*=0.0002) and remained elevated at 7 and 14 dpi (*p*=0.0008 and 0.0036, respectively), before returning to baseline at 35 dpi (Fig 1D). In humans, a significant increase in neopterin was detected between uninfected individuals and 1-14 DPO of ZIKV (*p*<0.0001; Fig 1E), indicating an upregulation of neopterin within the first two weeks of ZIKV infection, in agreement with our NHP results. Interestingly, the magnitude of neopterin formation was significantly higher in the CSF compared to the plasma at 7 dpi (p=0.0078 for both PTM and RM), possibly indicating increased localized inflammatory responses (Fig 1F).

In order to assess the overall level of peripheral inflammation within the first week of ZIKV infection, we characterized plasma levels of C-reactive protein (CRP), an acute-phase protein that has been used as an indicator of inflammatory responses (37). We observed that plasma levels of CRP increased by 4 dpi as compared to baseline, although this difference did not reach statistical significance (*p*=0.0645; Fig 1G). Taken together, these data indicate that ZIKV infection of PTM, RM and humans results in elevated systemic and CNS inflammation.

### Tryptophan catabolism is increased in plasma and CSF throughout ZIKV infection

Previous work has shown that tryptophan metabolism by indoleamine-2,3-dioxygenase (IDO) (17–20) is upregulated during viral infections (18,38), and significant changes to the kynurenine-tryptophan ratio (KTR) could be due to increased monocyte activation, and is associated with microbial dysbiosis (27). Moreover, increased amounts of kynurenine in the CNS have been linked to cognitive disorders and neurological deficits (39). We therefore measured levels of kynurenine (K) and tryptophan (T) in the CSF of ZIKV infected PTMs and RMs. The KTR was calculated, with a higher ratio indicating increased tryptophan catabolism. We detected a significant increase in the KTR in the CSF of PTM by 7 dpi, and this increase was sustained until 14 dpi (p=0.0056 and 0.0021, respectively) before returning to baseline at 21 dpi (Fig 2A). Conversely, no differences in the KTR in the CSF of RM was observed at any post-ZIKV infection time point as compared to baseline (Fig 2B).

**Figure 2.**
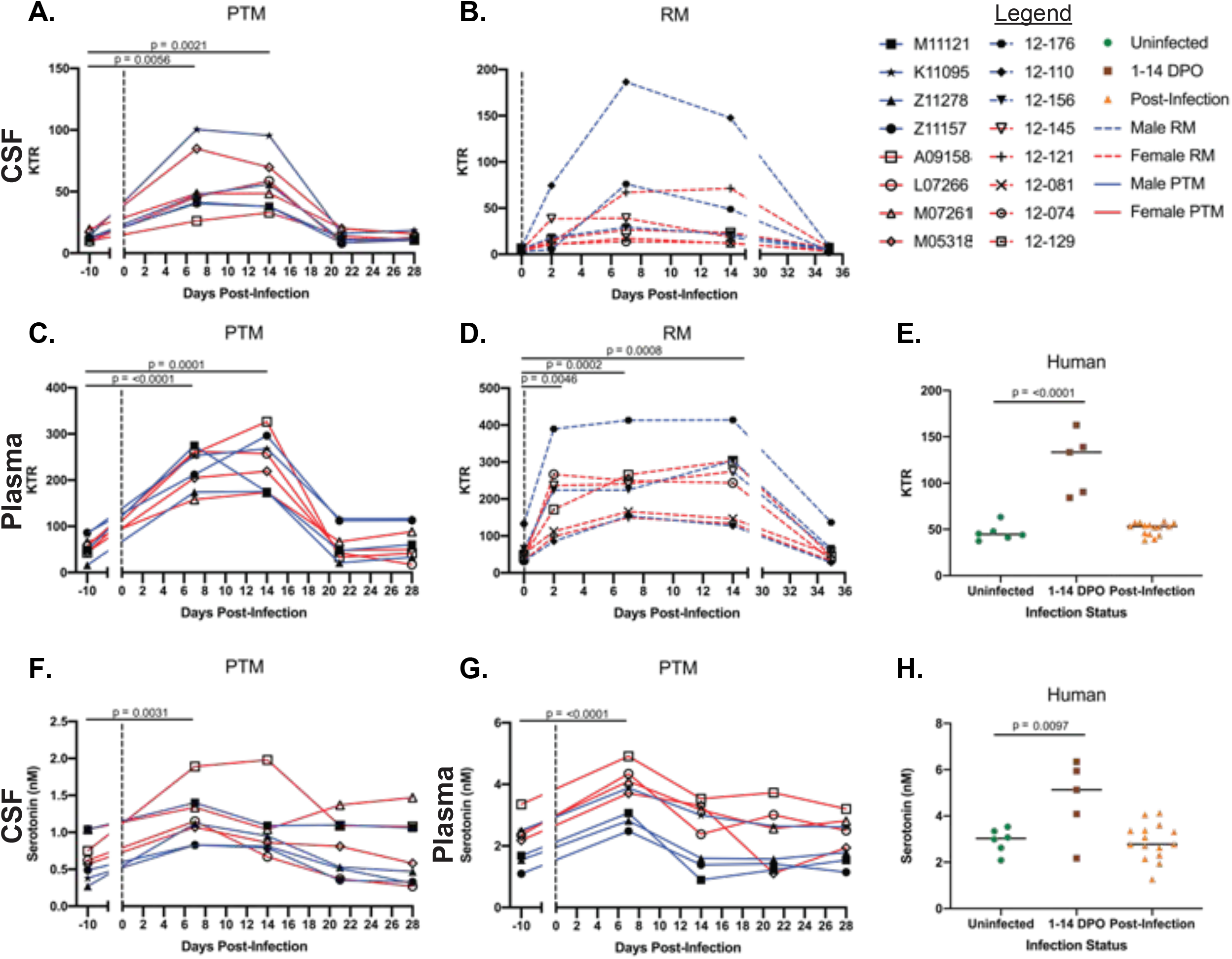
Increased kynurenine:tryptophan ratio (KTR) and serotonin levels are during ZIKV infection. Liquid chromatography tandem mass spectrometry (LC-MS/MS) was used to detect kynurenine and tryptophan concentrations in the CSF of PTM and RM and in the plasma of PTM, RM and humans throughout ZIKV infection. The KTR was calculated and the quotient was multiplied by 1000. Additionally, serotonin was measured in the plasma and CSF of PTM during ZIKV infection by LC-MS/MS. (A-E) KTR in the CSF of PTM (A), CSF of RM (B), plasma of PTM (C), plasma of RM (D) and serum of humans (E) infected with ZIKV. (F-H) Serotonin concentrations in the CSF of PTM (F) plasma of PTM (G) and serum of humans (H) infected with ZIKV. In panels A-D and F-G, each animal is represented by a different symbol. Solid or dashed lines connect data from the same animal. Solid blue lines indicate male PTMs; solid red lines indicate female PTMs; dashed blue lines indicate male RMs, dashed red lines indicate female RMs. Vertical dotted lines indicate the time point at which animals were inoculated with ZIKV. Statistical significance between the pre-infection baseline and post-ZIKV infection time points was calculated using a repeated measures ANOVA, with the Geisser-Greenhouse correction and a Dunnett’s multiple comparisons post-test. In panels E and H, uninfected humans are indicated by green circles, humans experiencing ZIKV symptoms from 1-14 days post-onset (DPO) of ZIKV symptoms are indicated by brown squares and humans post-ZIKV infection (15-156 DPO) are indicated by orange triangles. Black bars indicate median values. Statistical significance between human sample timepoints was evaluated using a one-way ANOVA with a Dunnett’s multiple comparisons post-test. In all plots, horizontal bars with p-values indicate time points that were statistically significant from each other.

To determine if the kynurenine pathway of tryptophan metabolism is upregulated in the periphery during ZIKV infection, we assessed the KTR in plasma of ZIKV infected PTMs, RMs and humans. A significant increase in the KTR was detected in the plasma of the PTMs 7 dpi (*p*=<0.0001) and sustained until 14 dpi (*p*=0.0001), returning to baseline by 21 dpi (Fig 2C). Similarly, a significant increase in the KTR was observed in the plasma of the RMs as early as 2 dpi (*p*=0.0046) and remained significant at 7 and 14 dpi (p=0.0002 and 0.0008, respectively), before returning to baseline at 35 dpi (Fig 2D). Additionally, a significant increase in KTR was observed in ZIKV-infected humans at 1-14 DPO of symptoms compared to uninfected individuals (*p*=<0.0001; Fig 2E), in agreement with our NHP results.

While production of kynurenine accounts for approximately 90% of tryptophan metabolism, approximately 3% is used for the synthesis of serotonin (5-HT) (40). Given our findings that ZIKV infection results in alteration of the KTR, we next investigated whether serotonin levels differed in the plasma and CSF of PTMs and in the plasma of humans. Serotonin levels were not measured in RMs due to lack of sample availability. A significant increase in serotonin levels was detected in the CSF of PTMs by 7 dpi (*p*=0.0031) and was elevated at 14 dpi, although this increase was not statistically significant (Fig 2F). Serotonin levels in the CSF of PTMs returned to baseline by 21 dpi (Fig 2F). Further, serotonin levels in the plasma of PTMs significantly increased 7 dpi (p=<0.0001) and returned to baseline by 14 dpi (Fig 2G). In human plasma, a significant increase in serotonin levels was seen between uninfected individuals and individuals at 1-14 DPO with ZIKV infection (*p*=0.0097; Fig 2H). Together, these data reveal that both tryptophan metabolism pathways become over-activated during the first week of ZIKV infection, which could be a possible mechanism involved in ZIKV pathogenesis and neurological complications that occur due to infection.

### Microbial translocation may contribute to elevated sCD14 in CSF during ZIKV infection

Increased microbial translocation has previously been observed in many viral infections, and are consistently associated with systemic inflammation and co-morbidities(28,41–45). We hypothesized that microbial translocation during ZIKV infection could contribute to increased monocyte activation and result in inflammatory responses. Cluster of differentiation 14 (CD14) is co-receptor for the bacterial product LPS; it is expressed either as a glycosylphosphatidylinositol (GPI)-anchored membrane protein on monocytes or macrophages (46) or as a soluble protein (sCD14) that is produced via secretion or enzymatic cleavage from the cell surface (47). sCD14 is considered a marker of monocyte activation due to microbial exposure (48), and translocated microbial products that enter the nervous system may induce sCD14 expression in this compartment (49). We thus measured sCD14 concentrations in the CSF throughout ZIKV infection of PTM and RM. A significant increase in sCD14 was detected in the CSF 7 dpi in the PTMs and was sustained through day 14 dpi (*p*=0.0124 and 0.0249, respectively; Fig 3A). In the RMs, a significant increase in sCD14 was detected in the CSF as early as 2 dpi (p=0.0253; Fig 3B), despite no detectable ZIKV RNA in the CSF at that time point (4). CSF sCD14 levels declined somewhat at 7 and 14 dpi in RMs but increased significantly at 35 dpi (*p*=0.0179; Fig 3B). We also examined sCD14 in the periphery but detected no significant changes in plasma sCD14 levels in either PTM or RM throughout ZIKV infection (Figs 3C and 3D, respectively). We also assessed serum CD14 levels in ZIKV infected humans and observed no significant differences in serum CD14 levels in uninfected individuals as compared to 1-14 days post-onset (DPO) and post-infection with ZIKV (Fig 3E).

**Figure 3.**
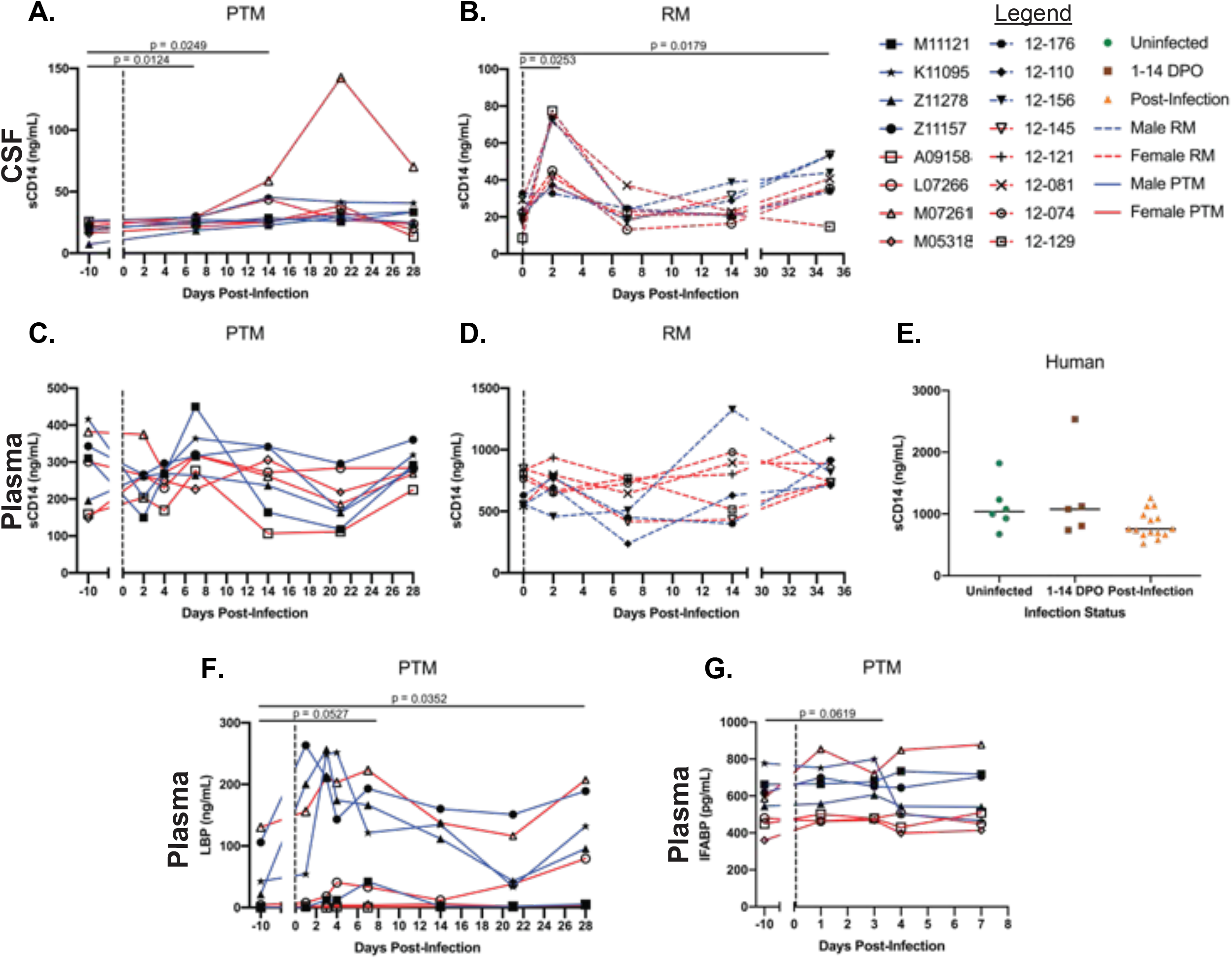
Increased levels of soluble markers of monocyte activation, microbial translocation and intestinal barrier dysfunction during ZIKV infection. Enzyme-linked immunosorbent assays (ELISAs) were used to detect soluble CD14 (sCD14), lipopolysaccharide binding protein (LBP) and intestinal fatty acid binding protein (I-FABP) in the CSF of PTM and RM and in the plasma of PTM, RM and humans throughout ZIKV infection. (A-D) sCD14 in the CSF of PTM (A), CSF of RM (B), plasma of PTM (C), plasma of RM (D) and serum of humans (E) infected with ZIKV. (F) LBP levels in the plasma of PTM. (G) I-FABP levels in the plasma of PTM. In panels A-D and F-G, each animal is represented by a different symbol. Solid or dashed lines connect data from the same animal. Solid blue lines indicate male PTMs; solid red lines indicate female PTMs; dashed blue lines indicate male RMs, dashed red lines indicate female RMs. Vertical dotted lines indicate the time point at which animals were inoculated with ZIKV. Statistical significance between the pre-infection baseline and post-ZIKV infection time points was calculated using a repeated measures ANOVA, with the Geisser-Greenhouse correction and a Dunnett’s multiple comparisons post-test. In panels E, uninfected humans are indicated by green circles, humans experiencing ZIKV symptoms from 1-14 days post-onset (DPO) of ZIKV symptoms are indicated by brown squares and humans post-ZIKV infection (15-156 DPO) are indicated by orange triangles. Black bars indicate median values. Statistical significance between human sample timepoints was evaluated using a one-way ANOVA with a Dunnett’s multiple comparisons post-test. In all plots, horizontal bars with p-values indicate time points that were statistically significant from each other.

Peripheral levels of lipopolysaccharide (LPS)-binding protein (LBP), an acute-phase protein that aids in binding of LPS to PRRs, has been used previously as a surrogate marker of microbial translocation (50–52). Thus, we assessed plasma levels of LBP before and after ZIKV infection of PTM. We observed a trend towards increased levels of LBP in plasma at 7 dpi (*p*=0.0527; Fig 3F). Plasma levels of LBP remained elevated and became significantly increased at 28 dpi (*p*=0.0352; Fig 3F).

Given the trend toward increased LBP in plasma and significant increases in sCD14 in the CSF, we next assessed for loss of mucosal epithelial barrier integrity. Intestinal fatty acid-binding protein (I-FABP) is a cytoplasmic protein found in intestine epithelial cells (4). I-FABP is released into the periphery after enterocyte disruption and thus has been used as a soluble biomarker of intestinal epithelial cell damage (53). To determine whether ZIKV infection caused intestinal epithelial barrier disruption, we evaluated plasma levels of I-FABP in ZIKV-infected PTMs during acute infection. Levels of I-FABP in the plasma during ZIKV infection remained consistent with the levels observed prior to ZIKV infection, although we did observe a trend towards increased I-FABP at 3 dpi (*p*=0.0619; Fig 3G), Taken together, our findings suggest that microbial product translocation into the periphery may contribute to increased CNS inflammation in ZIKV infection of PTM, RM and humans.

### Microbial taxonomy in the gastrointestinal tract of PTMs during ZIKV infection

Shifts in the GI microbiota occur frequently in the context of disease or infection (ie. Microbial dysbiosis), and such dysbiosis has been shown to alter mucosal integrity, peripheral PAMP levels, and local immune responses (27,28,54–58). West Nile Virus, a close relative of ZIKV, has previously been shown to promote significant GI alterations (59). To determine if ZIKV infection impacts the intestinal microbiome, we compared the relative abundance of rectal and stool microbiota of PTMs before and after ZIKV infection. Prior to infection, both rectal and stool microbiomes in male and female PTM were dominated by bacteria in the Firmicutes phyla, followed by Bacteriodetes and Proteobacteria (Figs 4A-D). An increase of Proteobacteria was detected 3 dpi in male rectal swabs, and levels remained elevated above baseline for the duration of the study (Figs 4A and 4E). This relative increase was reciprocated by a relative decrease in Firmicutes (Figs 4A and 4E). No shifts were detected post-infection at the phylum level in female rectal swabs (Figs 4C and 4E). Microbial phyla in male stool appeared relatively stable following infection (Figs 4B and 4E). Spirochaetes levels in female stool decreased at 3 dpi and remained below baseline throughout the study (Figs 4D and 4E). In addition to phyla level assessments, we also evaluated genus level changes in microbial communities throughout ZIKV infection. We observed a transient non-significant increase in *Prevotella* in stool from female PTMs at 3dpi, which decreased below baseline abundances by 28 dpi (S1 Fig). Additionally, there was a transient non-significant increase in *Streptococcus* in rectal swabs from female PTMs at 7 dpi, which returned to baseline abundances by 21 dpi (S1 Fig). No other shifts in microbial abundance were detected post-infection at the genus level in male and female rectal swabs and stool (S1 Fig).

**Figure 4.**
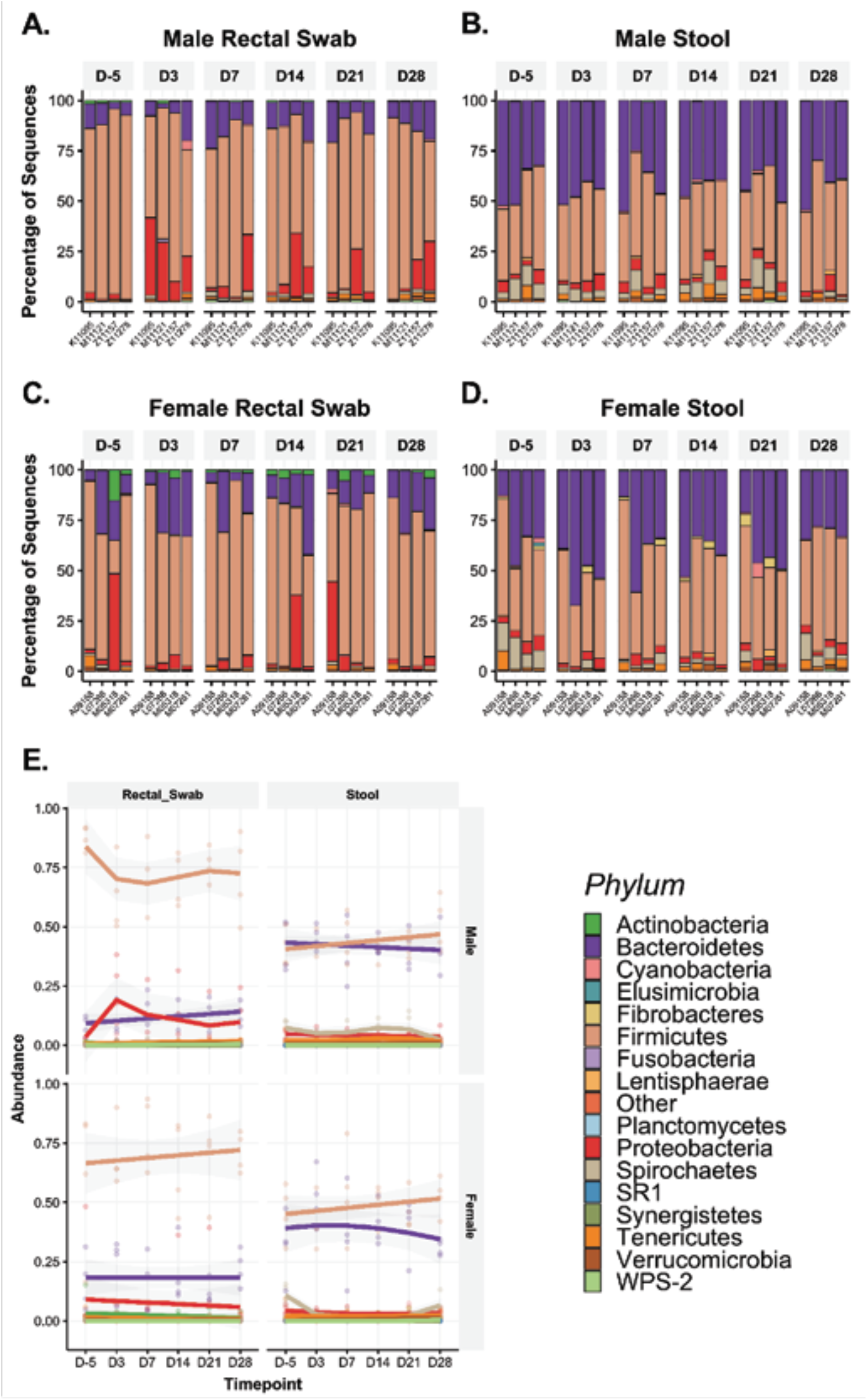
Bacterial community composition at the phyla level in the rectum and stool of PTMs before and after ZIKV infection. 16s rRNA gene sequencing was used to characterize microbial phyla in stool and rectal swabs collected from PTM prior to and throughout ZIKV infection. (A-D) Relative abundance taxonomic plots of microbial phyla in male PTM rectal swabs (A), male PTM stool (B), female PTM rectal swabs (C) and female PTM stool (D). Vertical colored bars represent the percentage of total sequences for specific phyla in individual animals prior to ZIKV infection and at each time point post-ZIKV infection. (E) Smoothed mean relative abundance of bacterial phyla in each of the indicated sample types in male and female PTM. Solid colored lines represent the mean abundance for specific bacterial phyla. Grey shading overlaying each colored line represents standard error bounds. Matched colored dots surrounding each colored line represent the specific abundances of each bacterial phyla for individual animals.

When considering the impact of ZIKV infection on microbiome structure, we did not observe significant changes in community richness or evenness (Figs 5A and 5B). Microbial diversity trends were not influenced by environmental variables such as origin of colony or paired caging (S1 Table), however, we did note a significant difference in microbiota richness between male and female PTMs in stool samples (S2 Fig). Analysis of microbiota composition as assessed through Bray-Curtis beta-diversity analysis demonstrated that microbial composition clustered by sex independent of infection status, suggesting that the strongest differences in overall microbial composition tracked with sex (Fig 5C). When segregated by sex, baseline samples fell within post-infection clusters for both rectal swabs and stool (Fig 5C). However, given that the male and female studies were performed at different times, cohort specific changes cannot be ruled out. We did note a small number of amplicon sequence variants (ASVs) that experienced statistically significant log_2_-fold shifts from baseline following infection (S3 Fig). However, these findings must be considered in the context of a single baseline observation, making it unclear if the changes observed were due to ZIKV-specific disease processes or were within the expected range of variation for a one-month span of time. In sum, these data indicate that ZIKV infection induced only subtle shifts in microbial taxonomy in the rectum and stool of male and female PTMs.

**Figure 5.**
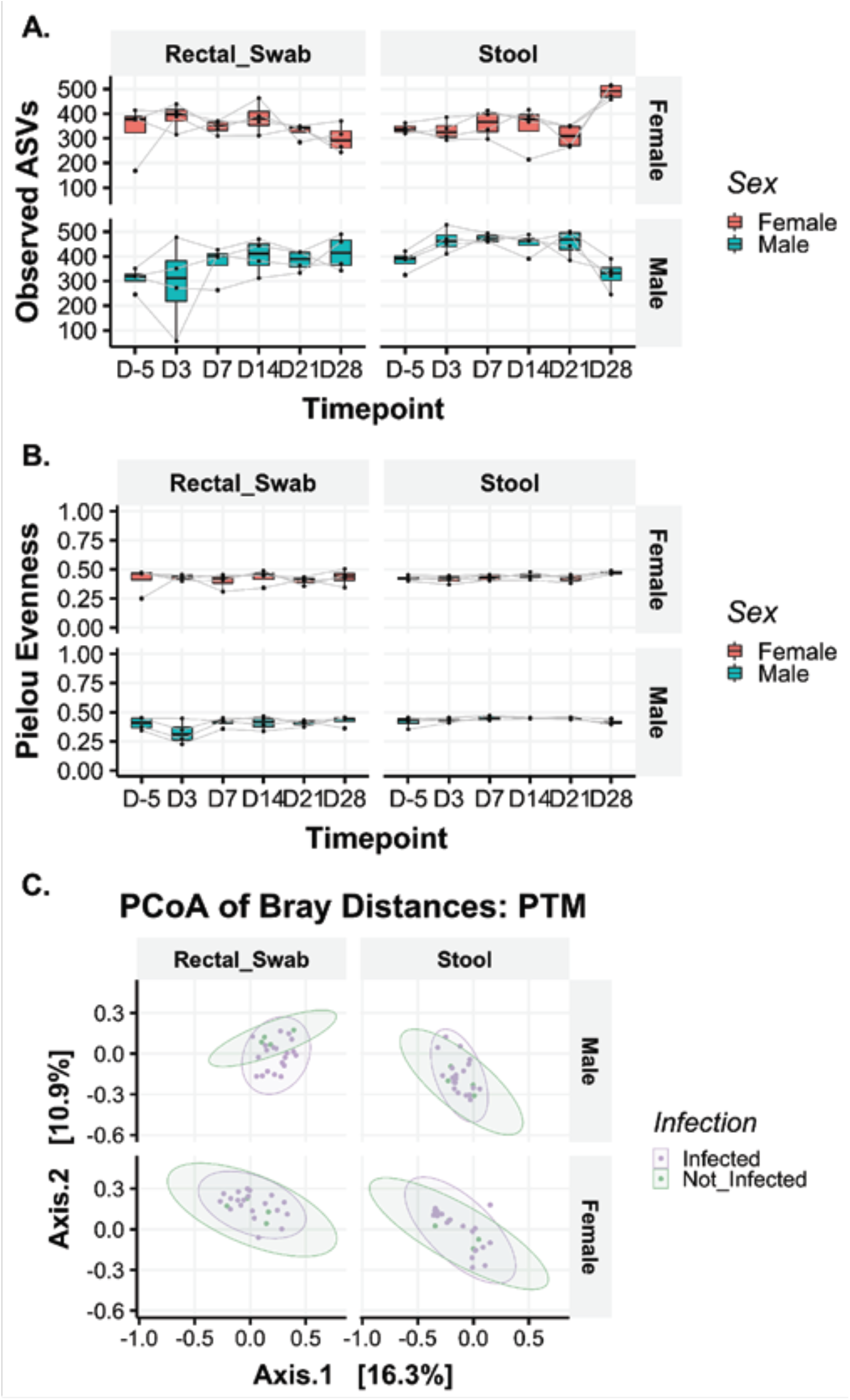
Similar bacterial community richness and evenness throughout ZIKV infection of PTMs. Measures of alpha-diversity were used to examine overall changes in stool and rectal bacterial community structure prior to and throughout ZIKV infection of PTMs. (A-B) Bacterial community richness (A) and evenness (B) in rectal swabs and stool from female (pink) and male (teal) PTMs. Box and whisker bars represent 25-75 percentile and minimum and maximum number of observed amplicon sequence variants. Horizontal bars within each box represent the median. Black dots connected by grey lines that overlay box and whisker plots represent the total number of observed ASVs for individual animals at each time point. (C) Principal components analysis of Bray-Curtis beta-diversity in male and female PTM rectal swabs and stool prior to and after ZIKV infection. Pre-ZIKV data points are shown in green and post-ZIKV data points are shown in purple. Shaded ovals for each group represent data ellipses.

### Depletion of short-chain fatty acids (SCFAs) in stool of PTMs during ZIKV infection

The intestinal microbiome contributes to GI health through multiple mechanisms, including by fermentation of metabolic products and production of short-chain fatty acids (SCFAs), which play a role in regulating immunity and inflammation (60,61). Although we did not observe significant alterations in microbial taxonomy in ZIKV infected PTMs, it is possible that subtle changes in community structure may still impact microbial functionality. Therefore, to evaluate intestinal microbial community function, we measured SCFA levels in the stool of ZIKV-infected PTMs. Acetate, butyrate and propionate, which are the primary end products of the fermentation process (62), were individually measured by GC-MS. Total SCFA production was also measured by GC-MS, which additionally includes measurements of iso-butyrate, valerate and iso-valerate. Concentrations of acetate in the stool pre-infection ranged from 0.70uM to 8.20uM and significantly decreased in all animals by 3 dpi (*p*=0.0018; Fig 6A). This decrease was sustained through 7 and 14 dpi (*p*=0.0026 and 0.0071, respectively) before returning to baseline at 21 dpi (Fig 6A). Butyrate concentrations pre-infection ranged from 1.33uM to 9.47uM and significantly decreased in all animals by 3 dpi (*p*=0.0343). As with acetate levels, this decrease was sustained through 7 and 14 dpi (*p*=0.0326 and 0.0573, respectively) before returning to baseline at 21 dpi (Fig 6B). Concentrations of propionate in the stool pre-infection ranged from 1.20uM to 14.28uM, and although levels were decreased by 7 dpi through 14 dpi as compared to baseline, this decrease did not reach statistical significance (*p*=0.0813 and 0.0534, respectively; Fig 6C). Levels of propionate returned to baseline at 21 dpi (Fig 6C). Additionally, total SCFA production significantly decreased at 3 dpi through 7 dpi (*p*=0.0232 and 0.0224, respectively; Fig 6D). Interestingly, female PTM began recovering total SCFA levels by 14 dpi, while male PTM sustained total SCFA depletion through 14 dpi (Fig 6D). Taken together, our findings indicate that although overall microbial community structure did not appear to be altered during ZIKV infection in PTMs, ZIKV may induce functional changes resulting in altered levels of microbial-derived metabolic products.

**Figure 6.**
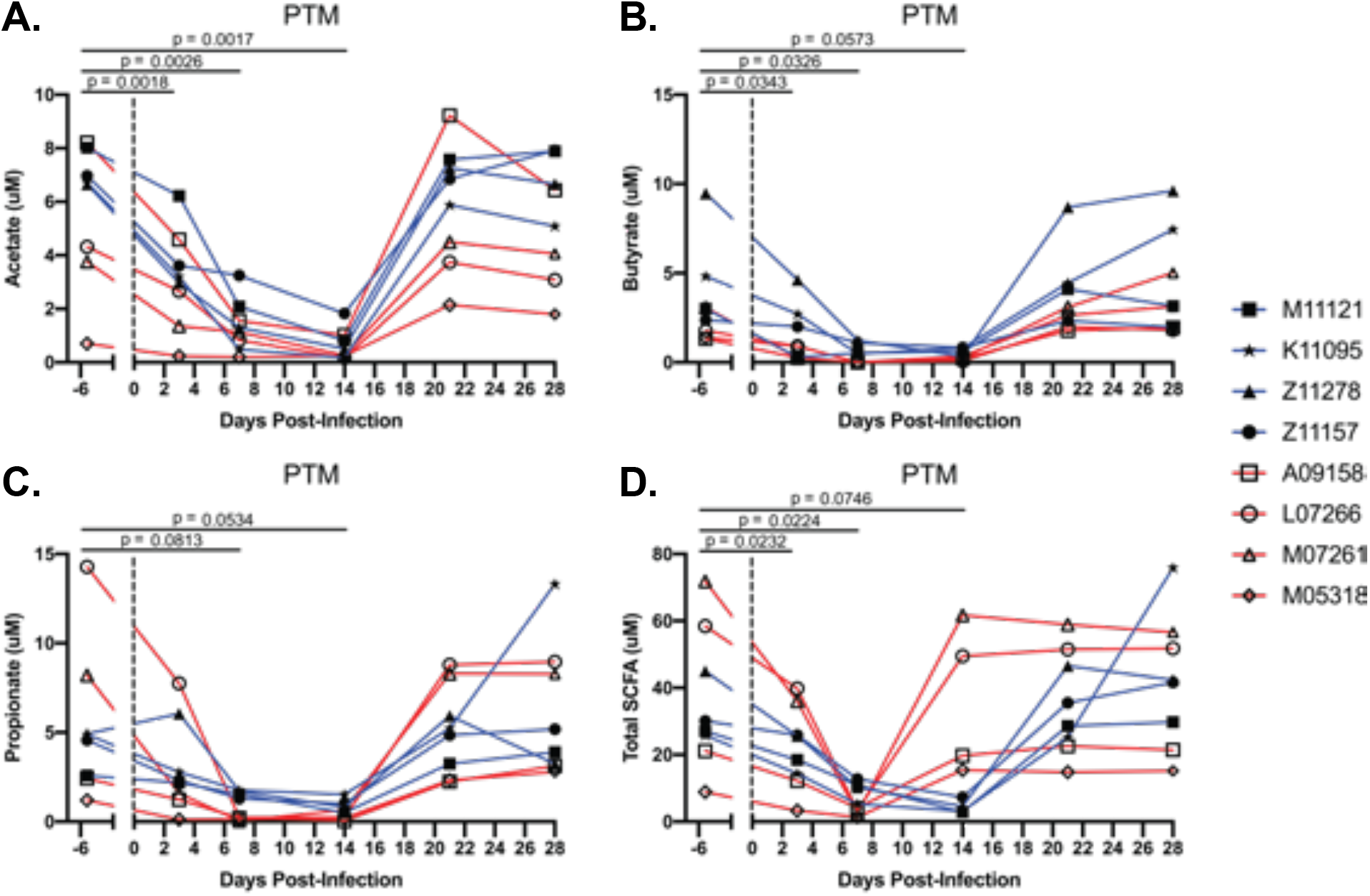
SCFAs are depleted in stool of PTMs during ZIKV infection. Gas chromatography mass spectrometry (GC-MS) was used to detect concentrations of short-chain fatty acids (SCFAs) in the stool of PTM throughout ZIKV infection. Concentrations of (A) acetate, (B) butyrate, (C) propionate, and (D) total short-chain fatty acids (SCFA) from stool were evaluated. Total SCFAs included concentrations of acetate, butyrate, propionate, iso-butyrate, valerate and iso-valerate. Each animal is represented by a different symbol. Solid lines connect data from the same animal. Solid blue lines indicate male PTMs; solid red lines indicate female PTMs. Vertical dotted lines indicate the time point at which animals were inoculated with ZIKV. Statistical significance between the pre-infection baseline and post-ZIKV infection time points was calculated using a repeated measures ANOVA, with the Geisser-Greenhouse correction and a Dunnett’s multiple comparisons post-test. In all plots, horizontal bars with p-values indicate time points that were statistically significant from each other.

### Associations between increased inflammatory biomarkers and indicators of microbial functionality during ZIKV infection

Previous studies have suggested that altered SCFA production may provide a mechanism by which microbial disruptions could influence peripheral and CNS inflammatory pathways production (63–67). To investigate if changes in SCFA levels were associated with changes in immunological parameters in ZIKV-infected PTM, we performed Spearman’s rank-order correlations between SCFAs and markers of inflammation in plasma and CSF (Fig 7). Analyses were separated based on male or female PTM study groups and data from all timepoints were used. We observed that in female PTMs, SCFAs were negatively correlated with plasma and CSF KTR (Fig 7A). These negative correlations reached statistical significance between the SCFAs acetate and butyrate and KTR in both the plasma and CSF (Fig 7A). SCFAs were negatively associated with serotonin levels, although this only reached statistical significance in the plasma with butyrate (Fig 7A). Similarly, SCFAs were negatively associated with neopterin levels, although this only reached statistical significance in the plasma with propionate (Fig 7A). Weaker negative associations were observed between all SCFAs and plasma and CSF sCD14 and plasma LBP (Fig 7A).

**Figure 7.**
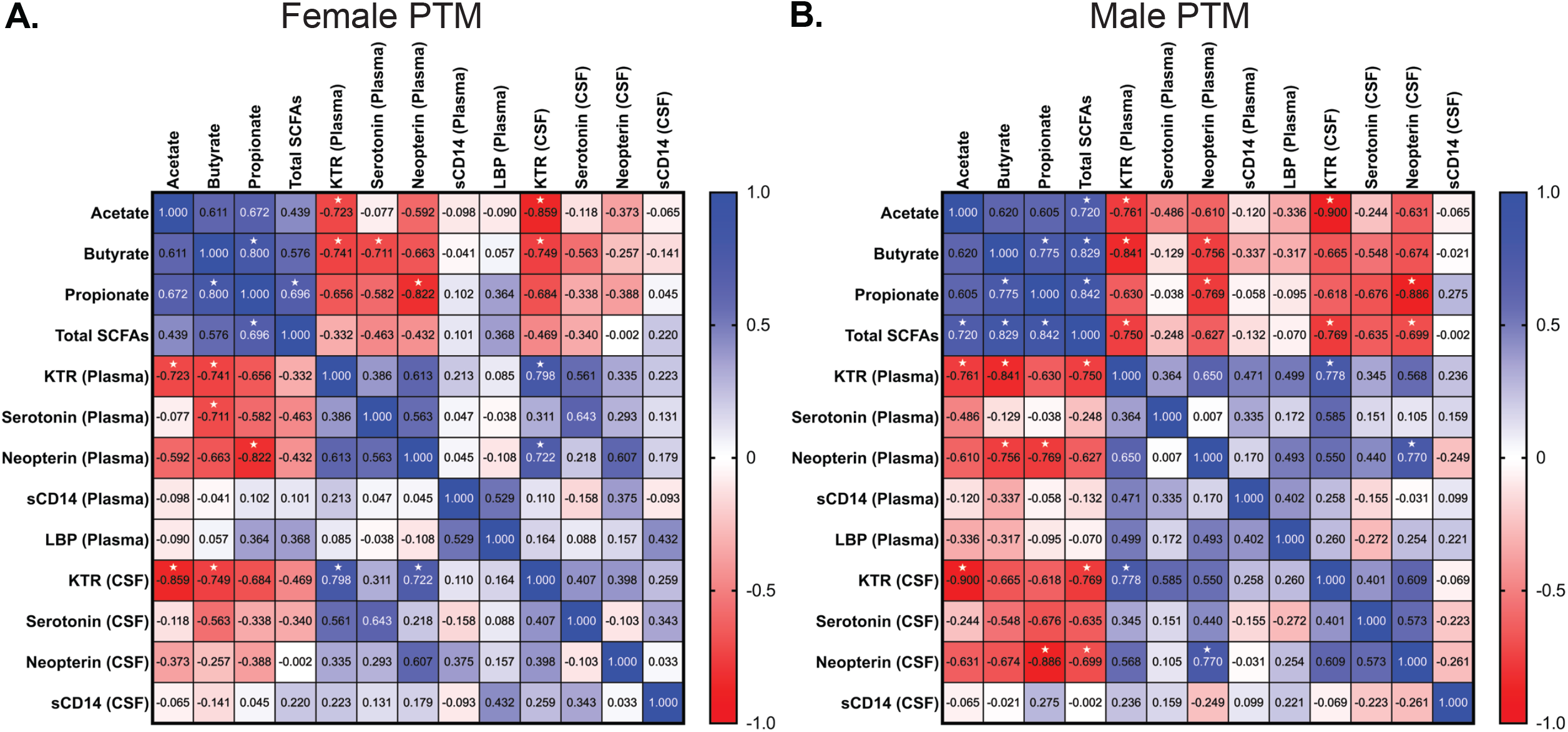
Decreased levels of stool SCFAs negatively correlate with increased plasma and CNS markers of inflammation and immune activation. Stool SCFA levels, plasma and CSF KTR, serotonin, neopterin, sCD14 and LBP concentrations in ZIKV infected PTM were correlated using a Spearman’s correlation. (A) Correlation matrix of stool, plasma and CSF markers in female PTM. (B) Correlation matrix of stool, plasma and CSF markers in male PTM. Data from all pre- and post-ZIKV infection time points for all analytes were used. Colors indicate Spearman correlation coefficient (R_s_) values ranging from −1 (dark red) to 1 (dark blue). Specific R_s_ values are depicted in the corresponding box for each comparison. Correlations which reached statistical significance (p<0.05) are indicated with a white star.

Similar trends were observed in male PTMs (Fig 7B). Specifically, SCFAs were negatively correlated with plasma and CSF KTR, and these associations reached statistical significance between plasma KTR and acetate, butyrate and total SCFAs, and between CSF KTR and acetate and total SCFAs (Fig 7B). Plasma and CSF neopterin levels were also negatively correlated with SCFA levels (Fig 7B). These negative correlations reach statistical significance between plasma neopterin and butyrate and propionate and between CSF neopterin and propionate and total SCFAs (Fig 7B). Plasma and CSF serotonin levels negatively associated with SCFAs, although none of these correlations reached statistical significance (Fig 7B). Weaker negative correlations were observed between SCFAs and plasma and CSF sCD14 and plasma LBP (Fig 7B). Taken together, these findings demonstrate that disruption of microbial SCFA production is associated with elevated peripheral and CNS inflammation during ZIKV infection, suggesting that alterations in microbial function may play a role in ZIKV pathogenesis.

## Discussion

The importance of the intestinal mucosal immune system and microbiome in maintaining health and homeostasis is widely recognized. However, the role of the mucosal immune system and microbiome in ZIKV infection, which recently became a rapid global epidemic (65), has not been well described. Indeed, given the significant associations between proper GI function and CNS homeostasis, it is possible that ZIKV-induced alterations in mucosal function could influence ZIKV pathogenesis in the CNS. Here, we used two separate NHP models, as well as clinical samples from ZIKV-infected humans, to explore whether ZIKV-induced elevations in systemic and CNS inflammation could be linked with increased mucosal dysfunction. We observed that ZIKV infection resulted in moderate elevations of microbial translocation markers in the plasma of PTMs, but significant increases in the CSF of PTMs and RMs. Of note, the three male PTMs with increased plasma LBP were the only animals that had detectable virus in the rectum at 7 dpi (24), suggesting a potential link between productive viral replication in rectal mucosal tissue and elevated intestinal barrier disruption, thus allowing for increased microbial translocation. Interestingly, we did not detect differences in plasma levels of the soluble marker of intestinal barrier damage, I-FABP, in ZIKV infected PTMs. However, given the notably short half-life of FABPs in the circulation (approximately 11 minutes) (68), it is possible that our sample collections time points were insufficient to capture transient changes in I-FABP levels. Further work assessing additional indicators of intestinal barrier damage will be needed to fully elucidate the impact of ZIKV on the mucosal epithelium and determine the potential mechanism underlying the loss of intestinal barrier integrity that may allow for elevated microbial translocation during ZIKV infection.

Our study revealed a significant increase in neopterin concentrations in the plasma and CSF of both species of NHPs after ZIKV infection. Interestingly, the concentration of neopterin was significantly higher in the CSF compared to plasma in both NHP species, potentially indicating a localized inflammatory response in the CNS during ZIKV infection. Moreover, our observation of elevated CNS levels of neopterin could indicate that activated monocytes are infiltrating and accumulating in the CNS, thus contributing to CNS pathology during ZIKV infection, as has been suggested in HIV infection (69). Our findings in the NHP model were in agreement with observations made using clinical samples, with significantly higher concentrations of neopterin found in the plasma of ZIKV-infected humans. Although we were unable to obtain CSF from ZIKV-infected humans, CSF neopterin concentrations could provide a biomarker or preliminary evidence of CNS inflammation and neurological symptoms in patients infected with ZIKV. Previous work has suggested that plasma neopterin levels are elevated in women with fetal growth restriction (70). It is possible therefore, that ZIKV-induced increases in neopterin in pregnant women could contribute to the development of ZIKV congenital syndrome. Additional work will be needed to determine the precise mechanism by which elevated neopterin levels occur in the plasma and CNS during ZIKV infection, and what role this could play in the development of fetal neurological deficits during gestation.

Concurrent with elevated neopterin levels, our study revealed a significant increase in the KTR in the plasma and CSF of ZIKV-infected PTMs and RMs. These findings were in agreement with observations made using clinical samples, with a higher KTR found in the plasma of ZIKV-infected humans. In combination with our observations of elevated plasma and CSF neopterin levels, higher peripheral and CNS KTR in ZIKV infection may indicate elevations in monocyte/macrophage activation, which could drive ZIKV neuroinflammation, similarly to what has been observed in HIV neuropathogenesis (69). Additionally, elevated catabolism of tryptophan in the CNS of ZIKV-infected individuals could result in higher levels of quinolinic acid, an end product of tryptophan degradation which has been linked to development of AIDS dementia complex (71). Further studies are needed to determine how altered tryptophan metabolism during ZIKV infection could contribute to systemic and CNS inflammation and immune activation, thereby driving ZIKV pathogenesis.

In addition to upregulated tryptophan metabolism to kynurenine, our study demonstrated increased serotonin levels in the plasma of ZIKV-infected PTMs and humans, and in the CSF of PTMs. In the brain, serotonin modulates critical neurodevelopmental processes, plays a role in cognitive function and affects mood in humans (40,72,73). A previous murine study suggested that maternal inflammation during pregnancy resulted in overactivation of serotonin synthesis, which contributed to adverse fetal brain development (72). It is therefore possible that maternal inflammatory responses during ZIKV infection could cause elevated serotonin production, which may in turn impact fetal neurological development. Additional work will be needed to fully examine the mechanism by which ZIKV-induced inflammation and disruption of metabolic processes, such as tryptophan metabolism into kynurenine and serotonin, may contribute to ZIKV pathogenesis and potentially ZIKV congenital syndrome.

Our assessment of stool and rectal microbial communities by 16s rRNA gene sequencing suggested that intestinal microbial community structure, including richness and evenness, is relatively stable in PTM following ZIKV infection. Although our study did not reveal major significant shifts in microbial taxonomy, we did observe decreased levels of SCFAs in the stool of ZIKV-infected PTMs, suggesting a potential impact of ZIKV infection on intestinal bacterial functionality. SCFAs have distinct physiological roles, including maintenance of gut barrier function, and have been shown to act as an energy substrate for colonocytes (74). Individuals with conditions like inflammatory bowel disease, obesity, or viral infections like HIV, have been shown to experience reductions in SCFA levels (75–78), which may promote further disruption of mucosal homeostasis. Our data demonstrating that SCFA depletion occurred after ZIKV infection could indicate that ZIKV causes generalized mucosal dysfunction. This alteration in mucosal homeostasis could drive chronic peripheral and CNS inflammation, thereby potentiating ZIKV pathogenesis. Taken together, our work suggests that regardless of taxonomic differences, ZIKV infection may impact microbial functionality, which may provide a mechanism underlying mucosal, systemic and CNS inflammation.

Our study had several limitations. First, the infections and sampling for the male and female PTM cohorts were performed at different times, so observed sex differences may be driven by the variability associated with performing study timelines at separate times. However, we attempted to minimize sampling bias further by generated all the data presented here together from cryopreserved specimens after sample collection for both cohorts were completed. Second, a limitation of our microbiome analysis is that our baseline sampling included only one pre-infection time point. This limits our ability to establish whether any observed community shifts were due to normal variation or a true biological effect of ZIKV infection. Considering the impact that infectious disease processes have on microbial community composition, future assessments should include extensive baseline sampling to enable statistical assessments of variation pre- and post-infection. Third, a limitation of collecting SCFAs from stool is the volatility of the esters. There is a chance that not all SCFAs were captured during these methods, as these samples were exposed to air for unknown amounts of time prior to collection. Finally, our work revealed some inter-animal variability in the inflammatory markers, microbial communities and bacterial-derived metabolites that were assessed. This variability is to be expected among outbred animals and could further be due to factors like primate origin, genetics and/or specific gut metabolism. The food provided was consistent between animals, thus diet is unlikely to be a factor in the subtle inter-animal differences observed.

In conclusion, our study explored mucosal dysfunction and microbial dysbiosis as a potential mechanism underlying ZIKV pathogenesis in two species of macaques, as well as in ZIKV-infected humans. Overall, our study suggests that ZIKV infection could result in decreased functionality of the gut microbiome, which could in turn result in decreased gastrointestinal health, increased mucosal inflammation and barrier damage. These disruptions could promote translocation of bacterial products into the periphery and contribute to the elevated inflammation observed in the periphery and CNS during ZIKV infection. Elevated inflammation may then induce upregulation of neopterin production and tryptophan catabolism in the CNS, which could contribute to the development of the neurological complications and deficits observed in patients infected with ZIKV. Additional studies are needed to better understand the mechanisms by which ZIKV and other viral infections such as SARS-CoV-2 infection may cause intestinal mucosal disruption, in particular alterations in the production of microbial-derived metabolites, and whether these disruptions could promote mother-to-fetal transmission of ZIKV and the development of fetal neurological defects.

## Materials and methods

### Ethics Statement

The pigtail macaques (*Macaca nemestrina*; n=8) used in this study were housed and cared for at the Washington National Primate Research Center (WaNPRC) under a protocol that was reviewed and approved by the University of Washington Office of Animal Welfare (OWA) Institutional Animal Care and Use Committee (IACUC; Protocol #201600397; Animal Welfare Assurance Number D16-00292). Animal housing, care and procedures were performed in an AAALAC-accredited facility, in accordance with the regulations put forth by the United States Department of Agriculture, including the Animal Welfare Act (9 CFR) and the Animal Care Policy Manual, and with the guidelines established by the National Research Council in the Guide for the Care and Use of Laboratory Animals and the Weatherall Report. Animals were housed in stainless steel cages with a 12/12 light cycle. All cage pans and animal rooms were cleaned daily and sanitized at least once every two weeks. Animals were provided with a commercial primate chow (Lab Diet, PMI Nutrition International) twice a day, with daily fruits and vegetables and water *ad libitum*. Male pigtail macaques were kept in run-through, paired housing throughout the entire study (M11121/Z11278; K11095/Z11157), while the females were in individual cages the entire study. Environmental enrichment consisted of novel food items, foraging opportunities and destructible and indestructible manipulanda. For minor procedures (Zika inoculation, blood and rectal swab collection) animals were anesthetized with ketamine (10mg/kg) and dexmedotomidine (0.017mg/kg). Of note, on day seven of the study, female pigtail macaques had ZIKV-associated weight loss which resulted in an over draw of blood by 1-3 mL in these four animals relative to IACUC guidelines of 10 ml/kg/week. For more involved sample collection, including CSF collection, general anesthesia was maintained with isoflurane by inhalation and post-operative analgesia was provided. Euthanasia was performed via an IV overdose (>75mg/kg) of pentobarbital in accordance with the recommendations in the Guidelines for the Euthanasia of Animals set forth by the Panel on Euthanasia of the American Veterinary Medical Association (AVMA).

Existing plasma and CSF samples from Rhesus macaques (*Macaca mulatta*; n=8) collected in a previous study (4) conducted by Dr. D.H. Barouch at the Center for Virology and Vaccine Research at Beth Israel Deaconess Medical Center were used in the present study. Briefly, rhesus macaques were housed and cared for at Bioqual, Rockville, MD under a protocol approved by Bioqual and the IACUC of Beth Israel Deaconess Medical Center, Harvard Medical School. As previously described, all animals were housed in single cages throughout the study and had not been previously enrolled in any other prior studies.

### NHP sample collection and processing

As previously described (24), eight healthy pigtail macaques (Cohort 1: 4 non-pregnant females, 8-10 years, 6-8 kg; Cohort 2: 4 males, 4-6 years, 10-12 kg) were subcutaneously infected with 5×10^5^ plaque forming units (PFU) of a Brazilian Fortaleza isolate of Zika virus (Brazil_2015_MG, GenBank: KX811222.1). Working stocks of ZIKV for the infection inoculum were grown and plaque-purified, as previously described (24). All animals were pre-screened for presence of antibodies to West Nile, Dengue, and Zika viruses, and all were negative except a single female animal (L07226), who was seropositive for West Nile. Antibiotics were not administered throughout the entire study timeline and were not administered within a year of study enrollment according to WaNPRC Animal History Reports. Diarrhea was not recorded in the Animal History Reports for any of the pigtail macaques during the time of this study. Study time points were referred to as the number of days post-ZIKV inoculation (dpi). Although male and female cohorts were sampled at different times, all sedation/sampling protocols remained the same between groups. Cerebrospinal fluid (CSF) was collected 10 days prior to inoculation and on days 7, 14, 21 and 28 dpi. To do this, animals were fasted for 24hrs prior to sedation and CSF was collected and cryopreserved neat at −80C for later analysis. Non-invasive samples, including stool, rectal and vaginal swabs and peripheral blood were taken 10 and 4 days before ZIKV inoculation and at days 1-4, 7, 14, 21 and 28 dpi. Stool and rectal and vaginal swabs were cryopreserved at −80C for later analysis. Blood sample were collected via venipuncture into EDTA vacutainers (BD, Franklin Lakes, NJ). Plasma was separated from whole blood by density gradient centrifugation and cryopreserved neat at −80C for later analysis. PTM were euthanized and necropsied either 28 or 29 dpi.

As previously described (4), eight Indian-origin, healthy rhesus macaques (5 females: 4-5 years, 4-5 kg; 3 males: 4-5 years, 5-6 kg) were subcutaneously infected with 1×10^3^-1×10^6^ PFU of ZIKV-BR 2015 and sampled at day 0 (pre-inoculation), 2, 7, 14 and 35 post-ZIKV. Animals were tested for baseline flavivirus neutralizing antibodies. Antibiotic treatments were not administered to these animals and of record, no GI symptoms (diarrhea) occurred throughout the study. CSF and plasma were collected at day 0 before inoculation (baseline time point), 2, 7 14 and 35 dpi. CSF was cryopreserved neat; plasma was separated from EDTA whole blood by density gradient centrifugation.

### Human Samples

Lyophilized human serum samples were obtained from the World Reference Collection for Emerging Viruses and Arborviruses, Galveston National Laboratory at the University of Texas Medical Branch. All samples from the Global Virus Network (GVN) Zika sera bank were donated, de-identified samples from various scientists and institutions. IRB #95-111 protocol states, “the samples will be obtained from discarded material collected for clinical purposes,” and a “No” to the statement “Will the donors be identified?” Additionally, within the IRB we acknowledge that “the proposed research is limited to the use of discarded materials or retrospective chart review and there are no identifiers associating the specimens or chart information with the donors.” Metadata for these sera samples included age, gender, place of exposure, date of illness onset, signs and symptoms, as well as days post-onset (DPO) for each collection time point. Study subject ages ranged from 15 to 67 years old. Sex was majority female with only 2/20 being males. Diagnostic tests were performed to verify ZIKV infection, such as qPCR and CDC lab diagnostics. Samples were grouped into either 1-14 DPO or post-infection (15-156 DPO). 5 samples were grouped as 1-14 DPO, the remaining 15 samples were in the post-infection group. To reconstitute the lyophilized human serum, samples were resuspended in the original volume stated on the vial using Hyclone^TM^ phosphate buffered saline (PBS; GE Healthcare Life Sciences; Pittsburgh, PA). The volume of samples ranged from 250 uL to 500 uL. Samples were vortexed until dissolved into solution. Finally, un-lyophilized plasma samples collected from healthy donors at the University of Washington/Fred Hutch Center for AIDS Research group at Harborview Medical Center were used as uninfected controls.

### Plasma assays

Soluble factors were assessed via enzyme-linked immunosorbent assay (ELISA) using the following commercially available kits: for soluble CD14 (sCD14), a Quantikine ELISA human sCD14 immunoassay from R&D Systems Inc. (Minneapolis, MN); for lipopolysaccharide binding protein (LBP), an ELISA kit from Biometic (Brixen, Italy); for monkey C-reactive protein (CRP), an ELISA kit from Life Diagnostics (West Chester, PA). Assays were completed according to the manufacturer’s recommended protocols and all samples were assessed in duplicate. Samples were diluted for each assay as follows: sCD14, plasma samples were diluted 1:200 and CSF samples were diluted 1:5; LBP: plasma samples were diluted 1:2.67; CRP: plasma was diluted 1:1000. Data was collected using an iMark microplate reader from Bio-Rad (Hercules, CA).

### 16s rRNA sequencing

<u>(i) Genomic DNA extraction and sequencing.</u> Genomic DNA was extracted from rectal swab specimens using lysis buffer, chicken egg white lysozyme, SDS and an RNA/DNA All-prep kit: DNeasy Blood and Tissue kit (Qiagen, Valencia, CA). DNA for 16S rRNA sequencing was processed using the Earth Microbiome Project protocols (28,41–44) with the following modifications. During the library preparation, each DNA sample was amplified in triplicate using the FailSafe™ PCR System (Epicentre, WI) and the 515FB-806RB primer pair to generate a 400 bp amplicon from the V4 variable regions of the 16S rRNA gene. The triplicate reactions were pooled, quantified using Qubit dsDNA High Sensitivity Assay Kit (ThermoFisher Scientific, Waltham, MA), and visualized using a LabChip GX (PerkinElmer, MA). 10ng of each library was pooled. The pooled library was cleaned using MO BIO UltraClean® PCR Clean-Up Kit (MO BIO, Carlsbad, CA) and quantified using the KAPA Library Quantification Kit (KAPA Biosystems, Wilmington, MA). Sequencing was carried out as detailed in the EMP protocol; specifically, 7pM of the pooled library with 30% PhiX phage as a control was sequenced using a 300-cycle Illumina MiSeq Kit (Illumina, Inc., San Diego, CA), generating 150bp paired-end reads. Raw 16S sequencing reads were submitted to NCBI’s Sequence Read Archive, accession number: PRJNA966191.
<u>(ii) Analysis of gene sequence data.</u> Raw 16S rRNA gene sequence data was processed using the QIIME 2 software package (version 2018.2.0) via the DADA2 (79) pipeline option which performs quality filtering and identification of exact amplicon sequence variants (ASVs). We then used the Greengenes database (release gg-13-8-99-515-806-nb-classifier) for taxonomic assignments. ASVs that were unassigned at the Kingdom level were filtered out and we removed ASVs that were present in only one sample. We then performed principal-coordinate analysis (PCoA) on log-normalized counts and generated relative abundance plots as a percentage of counts of the top 20 genera in each tissue type using the phyloseq

R package 1.22 (80). To identify differences in taxa abundance relative to baseline, we used DESeq2 R package 1.18 (81) to fit taxa abundances into a negative binomial generalized linear model (GLM) with parametric fitting of dispersions to the mean and p-values calculated with Wald significance tests. Significant differences were defined as ASVs with thresholds of adjusted p-value < 0.05 and a greater than 1.5 log_2_-fold-change difference from baseline in at least one time point.

### Mass Spectrometry

<u>(i) Liquid chromatography tandem mass spectrometry (LC-MS/MS).</u> Tryptophan (L-Trp), kynurenine (L-Kyn), serotonin and neopterin (D-(+)-Neopterin) concentrations were determined in plasma and CSF by LC-MS/MS. This method was validated for its sensitivity, selectivity, accuracy, precision, matrix effects, recovery, and stability. 10uL of sample was spiked with 10uL of internal standard (IS; L-Trp-d5) and 10uL acidified mobile phase (AMP; 0.2%FA/0.05%TFA/1% acetonitrile (ACN; Millipore-Sigma) in water). Subsequently, 150uL of ice-cold methanol (MeOH; Millipore Sigma) was added and this mixture was allowed to rest for 30min at −20°C to support protein precipitation. After centrifugation (3000 x g, RT, 10min) supernatants were removed and evaporated to dryness under a gentle stream of nitrogen. Dry extracts were reconstituted in 40uL AMP. 20uL of sample was injected into the LC-MS/MS. Chromatographic separation was achieved using a gradient elution with a Chromolith Performance RP-C18 column (Millipore-Sigma). The column was maintained at 25°C throughout. Samples were subjected to positive electrospray ionization (ESI) and detected via multiple reaction monitoring (MRM) using a LC-MS/MS system (Agilent Technologies 6460 QQQ/MassHunter; Palo Alto, CA). Each transition was monitored with a 150-ms dwell time. Mobile phase A is 0.1% formic acid in H_2_O and mobile phase B is 0.1% formic acid in methanol. During pre-study validation, calibration curves were defined in multiple runs on the basis of triplicate assays of spiked samples and quality control (QC) samples. Calibration standards were prepared and ranged from 0.05-1000 nmol/L with an inter- and intra-day precision and accuracy of ≤10.1% with an r^2^ value of 0.9973±0.0037. The LOD for all analytes was 0.05 nmol/L with an LOQ of 0.1 nmol/L. Quantification was performed using MRM of the transitions of m/z 205.1→118.0, 209.1→94.1, 254.1→206.1, 177.1→160.0, 210.1→122.1 for tryptophan, kynurenine, neopterin, serotonin and the internal standard (IS) tryptophan-d5, respectively.
<u>(ii) Gas chromatography mass spectrometry (GC-MS).</u> Concentrations of six short chain fatty acids (SCFAs), including acetate (acetic acid), propionate (propionic acid), isobutyrate (isobutyric acid), butyrate (butyric acid), isovalerate (isovaleric acid) and valerate (valeric acid) were determined in stool by GC-MS. This method was validated for its sensitivity, selectivity, accuracy, precision, matrix effects, recovery, and stability. Total short-chain fatty acids included all six acids. 1mL of acidified water (pH∼2.5, 9% formic acid; Millipore-Sigma) was added per 100mg of stool sample. Samples were vortexed and centrifuged at max speed for 5min. 300uL of supernatant was removed and transferred to a fresh, sterile, microcentrifuge tube. 30uL of internal standard (IS; 2-methyl-valeric acid) was added to each sample. 300uL ethyl acetate was added to each sample, vortexed for 10mins and centrifuged at max speed for 5mins. 200uL was carefully removed from upper organic phase and transferred into glass insert and capped immediately with non-pre-slit cap. Samples were stored at −20°C until they were collected on the GC-MS in batches. The GC-MS system consisted of an Agilent 5973N MSD EI/CI with 6890 GC and autosampler (Agilent Technologies). Acquisition was done using Chemstation software (Hewlett-Packard, Palo Alto, CA, USA). The GC was fitted with a polyethylene glycol (PEG), fused silica capillary column Stabilwax/Carbowax (30m, 0.32mm id, 0.25um film thickness) and helium was used as the carrier gas at 1mL/min. An injection volume of 1uL with an injection temperature of 250°C was used. For every three fecal samples injected, a blank sample with diethyl ether was inserted to minimize carry-over effects. The column temperature was initially 100°C and held for 2min then increased to 200°C at 25°C/min and kept at this temperature for 2 min (8min total time). The detector was operated in electron impact ionization mode at an electron energy of 70 V, scanning 30-250 *m/z* range. The temperature of the ion source, quadrupole, and interface were 220°C, 140°C, and 270°C, respectively. During pre-study validation, calibration curves were defined in multiple runs on the basis of triplicate assays of spiked samples and QC samples. Calibration standards were prepared and ranged from 0.01-100 umol/L with an inter- and intra-day precision and accuracy of <7.4% with an r^2^ value of 0.998±0.0013. The LOD for all SCFA was 0.05uM. Quantification was performed using quantification of target ions of 60, 74, 88, 73, 87, and 73 for acetic, propionic, isobutyric, butyric, isovaleric, and valeric acid, respectively.

### Data and statistical analysis

Statistical analyses were performed using GraphPad Prism statistical software (version 7; GraphPad Software, San Diego, CA). For NHP studies, results obtained at the pre-infection time points (day-10 or day-5 depending on sample type) were compared to those obtained 7, 14, 21 and 27-29 dpi. The significance between pre-infection and multiple post-infection time points in PTM and RM were evaluated using a repeated measures ANOVA with the Geisser-Greenhouse correction and a Dunnett’s multiple comparisons post-test, where *p* values of <0.05 were considered significant. Statistical analysis of human samples was performed by comparing the uninfected control group to either ZIKV infected group (1-14 DPO and post-infection). Statistical significance was evaluated using an ANOVA with a Dunnett’s multiple comparisons post-test, where *p* values of <0.05 were considered significant. To assess correlations between SCFA levels and inflammatory markers, Spearman’s rank order correlations were performed and a Bonferroni correction applied for multiple comparisons (*p*<0.0006).

## Acknowledgements

We thank all veterinary staff of the Washington National Primate Research Center and Bioqual Inc. for conducting the animal studies. We would like to thank the following for donating the human ZIKV+ serum samples included in this manuscript: The Allergen Foundation, The Global Virus Network (GVN) Zika Task Force, specifically the Chairperson Scott Weaver, MS, PhD, Dionna Scharton, and the World Reference Center for Emerging Viruses and Arborviruses (WRCEVA) at the University of Texas Medical Branch (UTMB) – Galveston National Laboratory.

## Author Contributions

MAO, JTG, CJM, DHB, NRK, MGJ, and DHF designed and coordinated the PTM study. CJM and MAO led and processed samples from PTM study. DHB provided RM samples. EAS used extracted DNA to sequence 16s rRNA. ATG and CBD performed 16s rRNA analysis and statistics. RKC performed mass spectrometry assays and analyzed resulting data. CAE assisted in preparing and performing all experiments with CJM. THM and LS provided feedback on data interpretation and manuscript preparation. JS led clinical care of PTMs. CJM, JAM, ATG and NRK interpreted the results and wrote the paper with all co-authors.

The authors declare no competing interests.

## Supporting information

**Figure S1.**
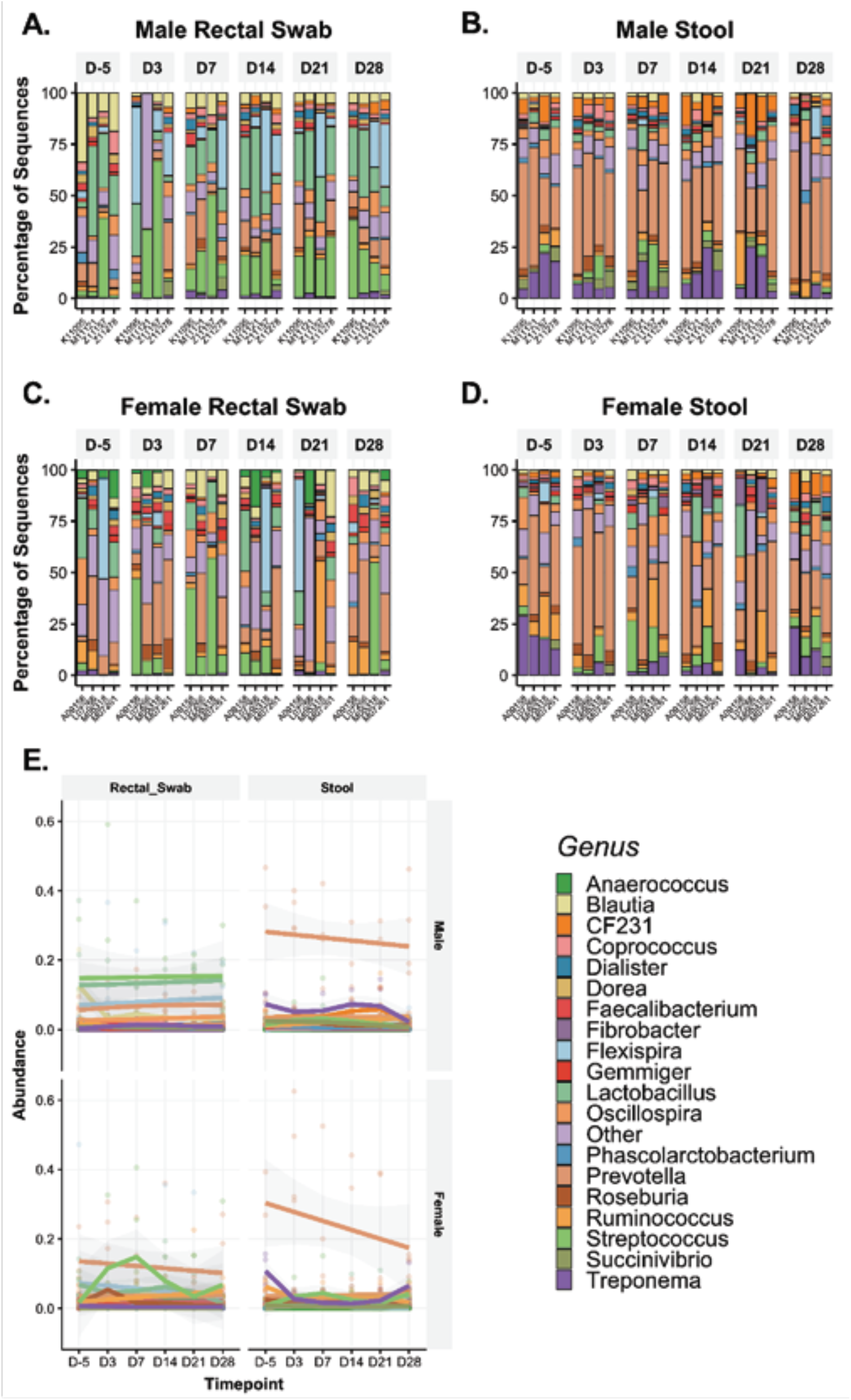
ZIKV infection induced minimal shifts in bacterial community composition at the genus level in the rectum and stool of PTMs. 16s rRNA gene sequencing was used to characterize microbial genera in stool and rectal swabs collected from PTM prior to and throughout ZIKV infection. (A-D) Relative abundance taxonomic plots of microbial genera in male PTM rectal swabs (A), male PTM stool (B), female PTM rectal swabs (C) and female PTM stool (D). Vertical colored bars represent the percentage of total sequences for specific genera in individual animals prior to ZIKV infection and at each time point post-ZIKV infection. (E) Smoothed mean relative abundance of bacterial genera in each of the indicated sample types in male and female PTM. Solid colored lines represent the median abundance for specific bacterial genera. Grey shading overlaying each colored line represents standard error bounds. Matched colored dots surrounding each colored line represent the specific abundances of each bacterial genera for individual animals.

**Figure S2.**
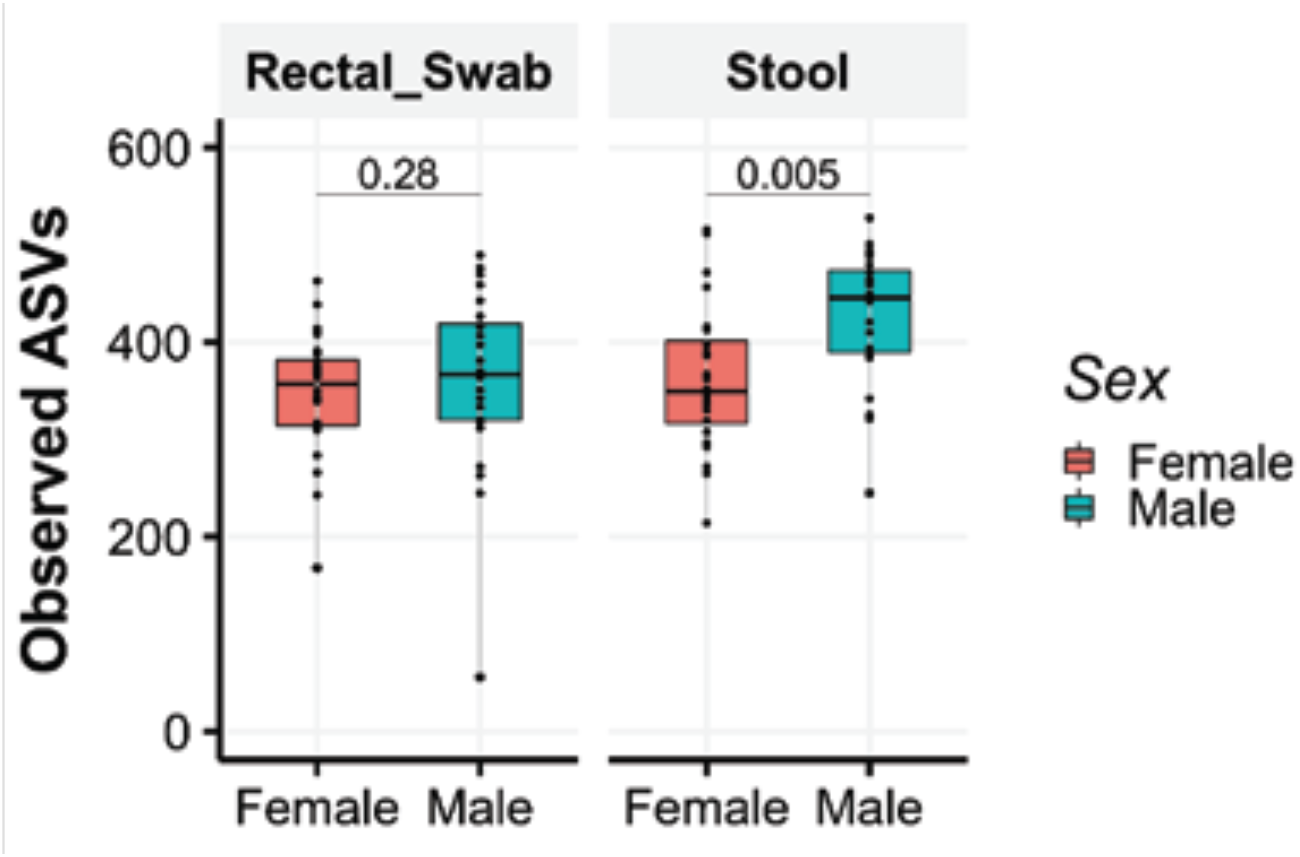
Increased richness in stool of male PTMs as compared to female PTMs. Bacterial community richness was assessed in rectal swabs and stool from female (pink) and male (teal) PTMs. Data from male and female PTMs at all time points were combined, regardless of ZIKV infection status. Box and whisker bars represent 25-75 percentile and minimum and maximum number of observed amplicon sequence variants. Horizontal bars within each box represent the median. Black dots that overlay box and whisker plots represent the total number of observed ASVs for individual animals at different time points. Statistical significance between the level of bacterial community richness in rectal swabs and stool of female and male PTMs was calculated using a Wilcoxon signed-rank test.

**Figure S3.**
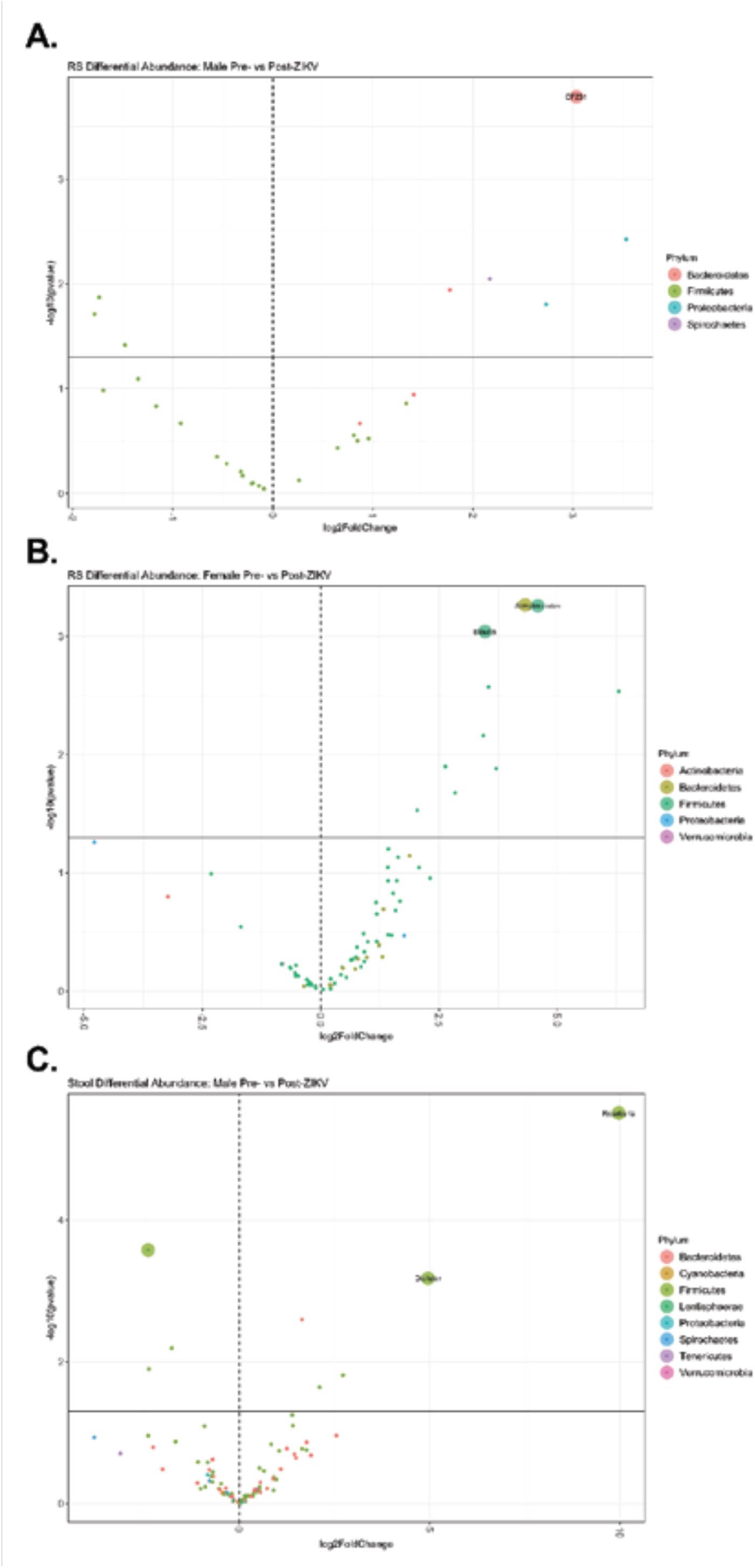
Alterations in the abundance of bacterial phyla following ZIKV infection. Shifts in bacterial ASVs away from baseline following ZIKV infection was assessed in stool and rectal swabs collected from male and female PTM prior to and throughout ZIKV infection. (A-C) Volcano plots depicting the log_2_-fold change in the abundance of ASVs after ZIKV infection in male PTM rectal swabs (A), female PTM rectal swabs (B), and male PTM stool (C). Colored dots represent individual bacterial ASVs. Large colored dots labeled with a genus indicate ASVs found to be significantly differentially abundant.

**Table S1.** Environmental variables for PTM.

| **Animal ID** | **Species** | **Origin** | **Sex** | **Age** | **Run-Through,**<br>**Paired Housing** |
| --- | --- | --- | --- | --- | --- |
| M07261 | PTM | WaNPRC at Tulane NPRC | F | 8 yrs | NA |
| A09158 | PTM | Yerkes NPC | F | 8 yrs | NA |
| L07266 | PTM | WaNPRC at Tulane NPRC | F | 8.5 yrs | NA |
| M05318 | PTM | WaNPRC at Tulane NPRC | F | 10.5 yrs | NA |
| M11121 | PTM | WaNPRC at SNBL TX | M | 5.5 yrs | Z11278 |
| Z11278 | PTM | WaNPRC at SNBL TX | M | 5.5 yrs | M11121 |
| K11095 | PTM | WaNPRC at SNBL TX | M | 5.5 yrs | Z11157 |
| Z11157 | PTM | WaNPRC at SNBL TX | M | 5.5 yrs | K11095 |

Potential environmental variables such as origin, sex, age and caging for the eight PTM.

